# Substrate Trapping in Polyketide Synthase Thioesterase Domains: Structural Basis for Macrolactone Formation

**DOI:** 10.1101/2024.06.20.599880

**Authors:** Tyler M. McCullough, Vishakha Choudhary, David L. Akey, Meredith A. Skiba, Steffen M. Bernard, Jeffrey D. Kittendorf, Jennifer J. Schmidt, David H. Sherman, Janet L. Smith

## Abstract

Emerging antibiotic resistance requires continual improvement in the arsenal of antimicrobial drugs, especially the critical macrolide antibiotics. Formation of the macrolactone scaffold of these polyketide natural products is catalyzed by a modular polyketide synthase (PKS) thioesterase (TE). The TE accepts a linear polyketide substrate from the termina PKS acyl carrier protein to generate an acyl-enzyme adduct that is resolved by attack of a substrate hydroxyl group to form the macrolactone. Our limited mechanistic understanding of TE selectivity for a substrate nucleophile and/or water has hampered development of TEs as biocatalysts that accommodate a variety of natural and non-natural substrates. To understand how TEs direct the substrate nucleophile for macrolactone formation, acyl-enzyme intermediates were trapped as stable amides by substituting the natural serine OH with an amino group. Incorporation of the unnatural amino acid, 1,3-diaminopropionic acid (DAP), was tested with five PKS TEs. DAP-modified TEs (TE_DAP_) from the pikromycin and erythromycin pathways were purified and tested with six full-length polyketide intermediates from three pathways. The erythromycin TE had permissive substrate selectivity, whereas the pikromycin TE was selective for its native hexaketide and heptaketide substrates. In a crystal structure of a native substrate trapped in pikromycin TE_DAP_, the linear heptaketide was curled in the active site with the nucleophilic hydroxyl group positioned 4 Å from the amide-enzyme linkage. The curled heptaketide displayed remarkable shape complementarity with the TE acyl cavity. The strikingly different shapes of acyl cavities in TEs of known structure, including those reported here for juvenimicin, tylosin and fluvirucin biosynthesis, provide new insights to facilitate TE engineering and optimization.

## Introduction

The number of pathogenic bacteria and fungi with evolved resistance to antimicrobial drugs is growing rapidly due to widespread clinical and agricultural use of antibiotics. This looming crisis places great urgency on the discovery and development of new pharmaceuticals. Macrolide antibiotics are an essential part of the antimicrobial arsenal, hence continual improvement in macrolide potency is essential to fight the ongoing war with pathogenic microbes. Exploiting natural biosynthetic systems, particularly for the challenging step of macrocycle formation, is an appealing approach to this problem^1^.

Bacterial type I modular polyketide synthases (mPKS) are nature’s biosynthetic machines for the macrolactone scaffold of macrolide antibiotics^2^ as well as for many anticancer drugs,^3^ fungicides^4^ and insecticides^5^. The mPKS biosynthetic pathways for macrolactones are organized as a sequence of multi-enzyme modules, each consisting of domains that carry, extend, modify and ultimately offload polyketides. An acyltransferase (AT), which may reside within or outside the module, selects a specific acyl-extender unit from the co-enzyme A (CoA) pool and transfers it to the phosphopantetheine (Ppant) prosthetic group of the module acyl carrier protein (ACP). Within the module, a ketosynthase (KS) catalyzes the decarboxylative Claisen condensation of the extender unit with the polyketide intermediate of the upstream module. Optional ketoreductase (KR), dehydratase (DH) and enoylreductase (ER) domains may modify the β-ketone to form a β-hydroxy, β-ene or fully reduced module product. Finally, at the terminus of the mPKS pathway, a thioesterase (TE) offloads the full-length polyketide product by cyclization to a macrolactone or hydrolysis to a linear carboxylic acid.

The formation of macrocycles by TEs during polyketide chain termination is of great importance to the development of biocatalysts and to other applications of pathway engineering. As catalysts of the final step in the PKS assembly line, TEs are essential for overall pathway flux and act as the final quality check prior to release of the polyketide product^6^. The macrolactonizing TEs form a wide range of natural products, and some can also cyclize non-natural polyketide substrates^7–9^. Synthetic approaches to lactonization of large substrates with numerous functional groups and chiral centers is generally challenging and low-yielding^10,11^.

Therefore, engineering TEs to broaden substrate scope or more finely control macrolactone production represents an exciting addition to the biocatalytic toolkit.

The basis for cyclization vs. hydrolysis by mPKS TEs (**Fig. 1A**) is poorly understood^6,12^ despite published structures of three macrolactone-forming TE domains, including the DEBSIII TE (erythromycin, 14-membered macrolactone 6-deoxyerythronolide B (6-deB))^13,14^, PikAIV TE (pikromycin, 12-membered 10-deoxymethynolide (10-dml) or 14-membered narbonolide)^15,16^, and PimS4 TE (pimaricin, 26-membered polyene ring)^17^; one hydrolyzing TE (TmcB TE, linear product leading to tautomycetin)^18^; and one decarboxylating TE (CurM TE, linear product curacin A)^19^. Biochemical study of DEBSIII TE and PikAIV TE revealed some substrate selectivity in macrolactone formation^8,9,15,20–23^. While tolerant of variation in substrate length and methyl or hydroxy substituents, DEBSIII TE was selective for the configuration of the nucleophilic alcohol^6,23^. Similarly, PikAIV TE tolerated certain changes in substrate length but required a C9 ketone to generate a macrolactone via the terminal hydroxy nucleophile at C13^15,20^ (**Fig. 1A**). Furthermore, cyclization by the PikAIV TE, like DEBSIII TE, is selective for one configuration of the nucleophilic hydroxyl^8^. These and other results^9^ implicate TEs as crucial quality-control gatekeepers for module processing of natural and unnatural substrates.

**Figure 1.**
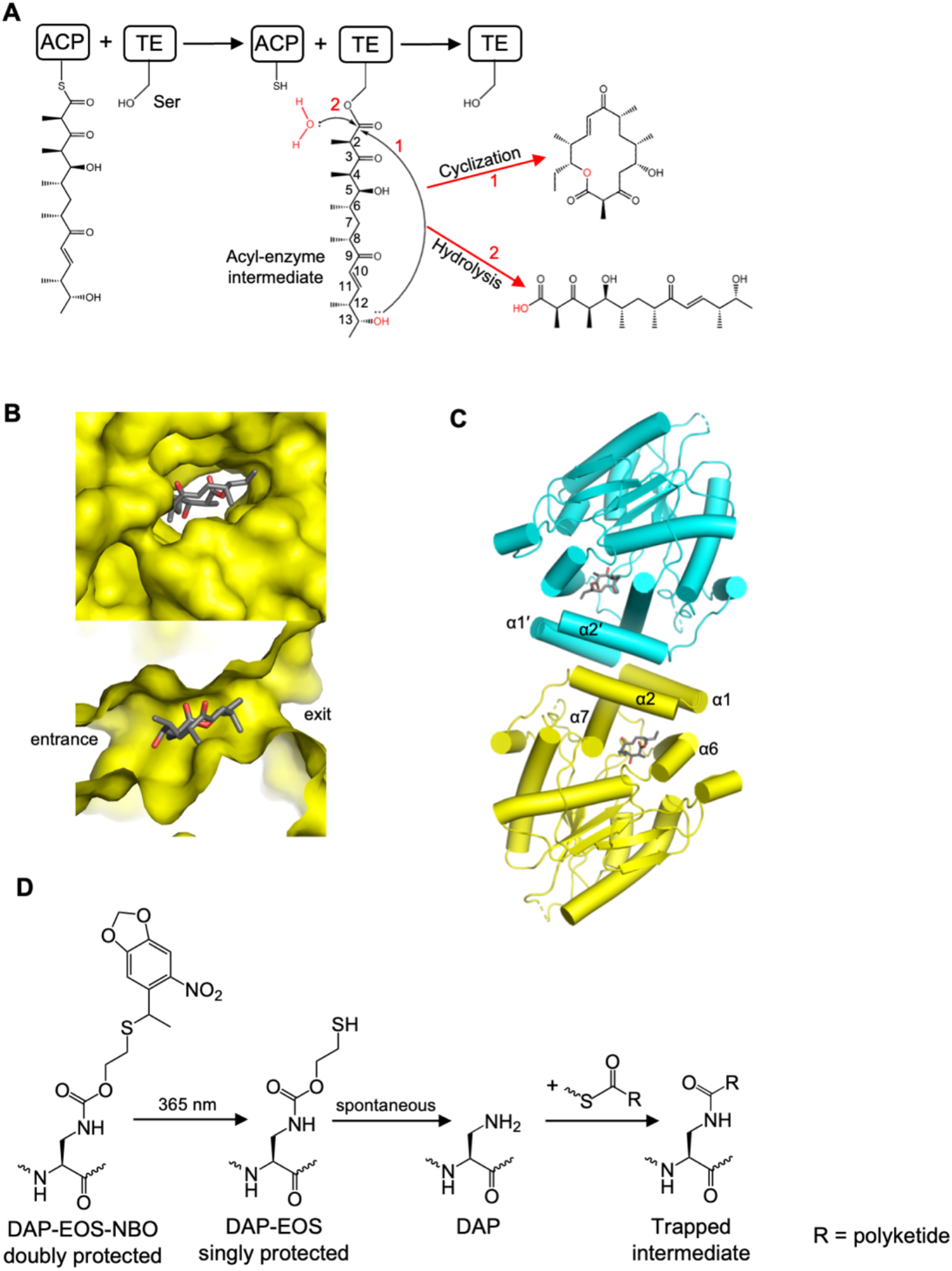
Macrolactone formation by offloading PKS thioesterases. **A**. Schematic of PikAIV TE offloading reaction with (1) macrolactone or (2) hydrolysis products. **B**. Perpendicular views of the active site tunnel in a PKS offloading TE dimer. The product 10-dml (stick form) is inside the acyl cavity in the surface rendering of the PikAIV TE (PDB 2HFK). The top image is viewed through the tunnel exit, and the bottom image is a cutaway from the side. **C**. Helices in the TE dimer interface. Helices α1-α2 form all dimer contacts and stabilize the active site tunnel through intra-subunit contacts with α6 and α7. Monomers are in contrasting colors. **D**. Unnatural amino acid deprotection scheme leading to DAP, followed by amide formation from a polyketide-thioester.

The offloading TEs in mPKS, nonribosomal peptide synthetase^24^ (NRPS) and fungal iterative PKS^25^ (iPKS) pathways are homologs within the α/β-hydrolase superfamily. They have remarkably similar structures even though they act on divergent substrates and have distinct macrolactone^13,15^, carboxylic acid^26^, macrolactam^27^, or decarboxylated^19^ products. The mPKS TEs are dimers with independent active sites. Within each monomer, a canonical catalytic triad (Ser or Cys nucleophile, His, Asp) is located at the center of a tunnel in which the cognate ACP can deliver the polyketide substrate through one end, and the offloaded product can exit through the opposite end (**Fig. 1B**). TE dimer formation is critical to the integrity of the active site tunnel. Two N-terminal α-helices (α1-α2) form all inter-subunit dimer contacts and stabilize the tunnel through intra-subunit contacts with helices α6-α7 (**Fig. 1C**). The tunnel architecture is unlike the monomeric TEs where a mobile lid, topologically equivalent to helices α6-α7, opens and closes over the active site, and no tunnel exists^28^. Helices α6-α7 have been described as a “lid” in dimeric PKS TEs^6^, but this lid is kept closed by helices α1-α2 in the dimer interface.

The exit end of the active site tunnel forms a spacious acyl cavity large enough to accommodate the polyketide substrate as well as the macrolactone or linear carboxylic acid product. The shape and extent of the acyl pocket (helices α2, α6, α7) is the most variable region in TE structures^13,15,17^, and includes several positions where sequences vary (**Fig. S1A**). No significant structural differences exist between mPKS TE free enzymes and their corresponding substrate, product or analog complexes^15,17,29^. Thus, substrate selectivity and catalytic outcome – macrolactone formation vs. hydrolysis – are not driven by conformational changes to the active site tunnel or acyl cavity but rather by other TE features such as the shape and chemical environment of the acyl cavity, the substrate structure, and the length, position and amphipathic profile of the lid region^6,13,15,17^. Visualization of active sites mimicking the covalent acyl-enzyme intermediate are key to a deeper understanding of the various parameters that control product outcome.

The intrinsic lability of the TE covalent intermediate – acyl-Ser ester or acyl-Cys thioester – has been a hurdle to direct visualization of TE-polyketide interactions. We previously used diphenyl- activated polyketide phosphonates to trap the PikAIV TE active site with phosphonate adducts that mimic the tetrahedral intermediate of acyl-enzyme formation^15,29^. However, the bulky diphenylphosphonate prevented full modification of the nucleophilic amino acid in the TE active site tunnel^15^. Here we adopted a recently developed platform to incorporate an amino functionality in place of the active site Ser hydroxyl or Cys thiol^30^. The unnatural amino acid (UAA) 2,3-diaminopropionic acid (DAP) is an isostere of Ser and Cys that can trap acyl-enzyme intermediates as stable amides (**Fig. 1D**). We tested DAP replacement for the nucleophilic amino acids in the offloading TE domains of five actinobacterial mPKS pathways (**Fig. S1B**).

Amide trapping was achieved for two TEs. A crystal structure of the PikAIV TE_DAP_ with the natural heptaketide illustrates how the structure of both the enzyme and the substrate contribute to the high fidelity of macrolactone formation. The acyl cavities of a panel of offloading PKS TEs, including three with structures reported here, reveal an overall correlation with substrate and product structures.

## Results

### Panel of PKS TE domains

The offloading TE domains of five actinobacterial type I PKS pathways were investigated for the possibility of replacing the nucleophilic Ser or Cys with the DAP isostere (**Figs. 1D, S1**): PikAIV TE (*Streptomyces venezuelae* ATCC 15439), DEBSIII TE (*Saccharopolyspora erythrea*), JuvEV TE (*Micromonospora chalcea* ssp. *izumensis*), TylG5 TE (*Streptomyces fradiae*) and FluC TE (*Actinomadura vulgaris*). We first purified all five wild-type TE domains (TE_WT_) to homogeneity and confirmed the expected dimeric state in solution (apparent molecular weight 55-65 kDa). The five TE domains are homologs with 29% – 45% pairwise sequence identity, except the JuvEV and TylG5 TEs, which produce the same tylactone product and have 64% identical sequences.

### DAP incorporation in PKS TE domains

DAP incorporation in our panel of five TEs followed by reaction with linear polyketide intermediates or macrolactone products provided a promising platform to probe the structural basis of macrolactone formation and substrate selectivity by the TEs. For DAP incorporation, the catalytic Ser or Cys codon in each TE-bearing plasmid was replaced with an amber stop codon, followed by co-expression with a plasmid encoding an evolved pyrrolysyl-tRNA synthetase (DAPRS) and a suppressor tRNA (tRNA^Amb^)^30^. DAPRS charges the tRNA^Amb^ with a DAP variant (doubly protected DAP-EOS-NBO, **Figs. 1D, S2**).

To optimize conditions for production of recombinant N-terminal His_6_-tagged TE_DAP_ in *E. coli* cells, we screened vector backbone (pMCSG7_31_ and derivatives pCDF7 and pTrc7), DAP-EOS- NBO concentration (1-10 mM), and culture temperature (18°C, 20°C, 25°C), and used SDS-PAGE to estimate the relative proportions of full-length TE (∼30 kDa) and truncated TE (∼16 kDa) resulting from non-incorporation of DAP-EOS-NBO. We identified conditions to produce useful quantities of PikAIV TEDAP and DEBSIII TE_DAP_ (**Fig. S3**) with yields 10-25 times lower than the corresponding TE_WT_ (9.1 mg/L vs. 100 mg/L for PikAIV TE; 7.2 mg/L vs. 130 mg/L for DEBSIII TE). No combination of conditions yielded sufficient quantities of TE_DAP_ for the JuvEV, FluC or TylG5 TEs, for which yields of the wild type proteins were far below the high levels for the PikAIV and DEBSIII TEs.

In forming TE_DAP_, the initial UAA was the doubly protected DAP-EOS-NBO (TE_DAP_+297.03 Da, **Fig. 1D**). Deprotection involved photo cleavage (365 nm) to the singly protected DAP-EOS (TE_DAP_+103.99 Da) followed by spontaneous deprotection to the active DAP (**Fig. S4**). The spontaneous deprotection step was slow for the PKS TE_DAP_ domains, perhaps due to slow solvent exchange in the substrate tunnel. We detected all three states of PikAIV and DEBSIII TE_DAP_ (TE_DAP-EOS-NBO_, TE_DAP-EOS_, TE_DAP_) by liquid chromatography / mass spectrometry (LC/MS) analysis of the intact protein in fresh preparations (**Fig. S4**). Complete photo deprotection occurred with UV exposure immediately following the Ni-affinity purification step. Conditions for the second spontaneous deprotection (temperature, time, pH, addition of reducing agent) were optimized to generate purified PikAIV TE_DAP_ and DEBSIII TE_DAP_ in the fully deprotected state.

### Covalent trapping of polyketides

We next characterized the reactivity of PikAIV and DEBSIII TE_DAP_ with linear polyketide substrates: thiophenol-activated forms of the pikromycin pentaketide^32^ and 3-hydroxymethyl-hexaketide^33^, and evaluated the formation of stable adducts using intact-protein LC/MS. For these linear substrates, PikAIV TE_DAP_ was fully modified with both adducts (**Figs. 2A, S5A,B**). DEBSIII TE_DAP_ was fully modified by the pikromycin pentaketide and modified at a lower level by the pikromycin hexaketide (**Figs. 2A, S6A,B**).

**Figure 2.**
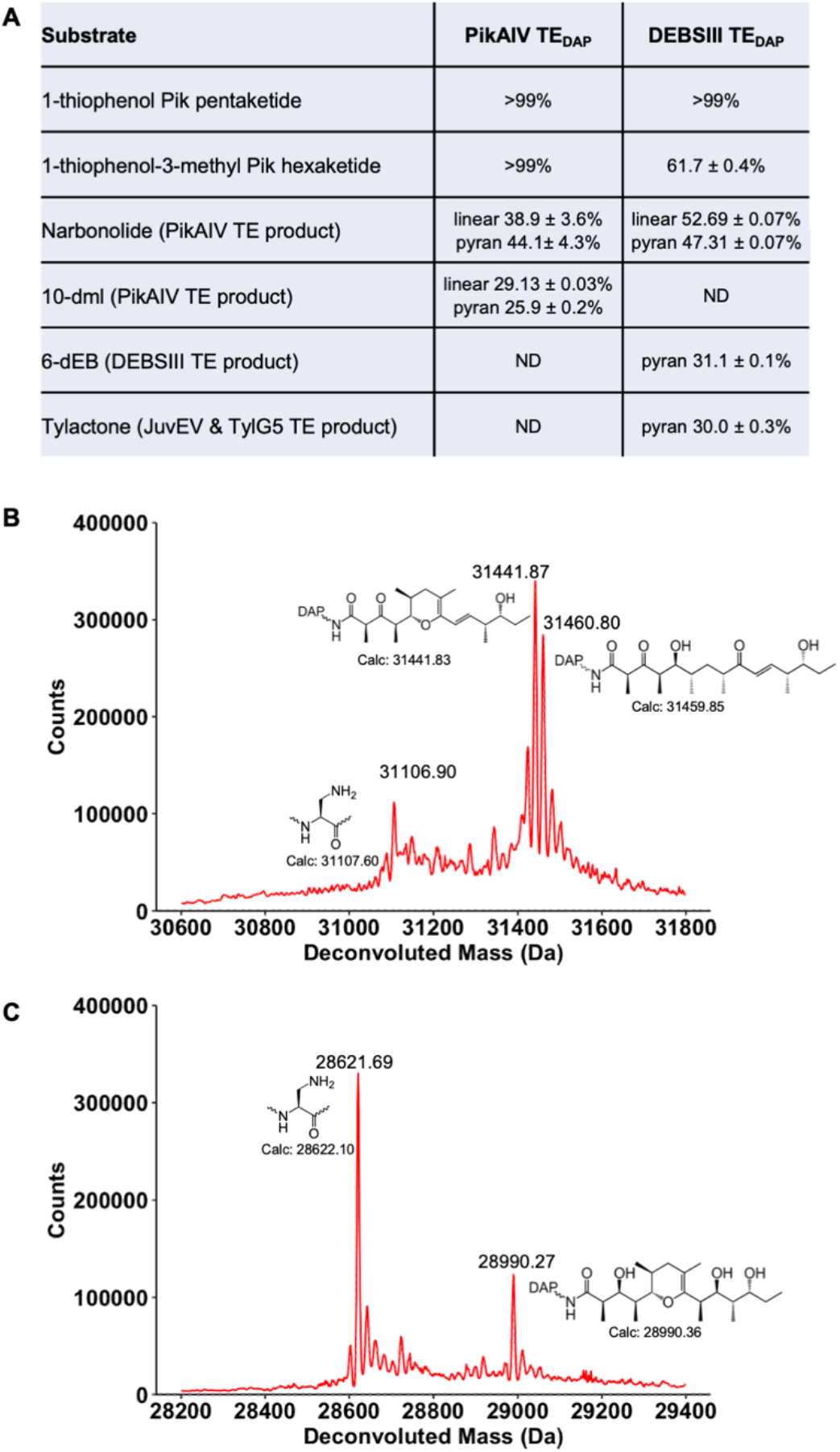
Polyketide modification of TE_DAP_. **A**. Quantitation of adduct formation for PikAIV and DEBSIII TE_DAP_ with a panel of linear and cyclic polyketides in a 24-hr reaction at room temperature. Values are based on peak areas in deconvoluted mass spectra for the intact proteins. Variances are based on triplicate measurements. **B**. PikAIV TE_DAP_ adducts from narbonolide. TE_DAP_ opened the 14-membered macrolactone ring, trapping the natural heptaketide as a stable, linear acyl-enzyme adduct. Deconvoluted mass spectra show the incorporation of heptaketide at DAP1196. The dehydrated pyran product resulting from spontaneous hemiketal formation also accumulated. **C**. DEBSIII TE_DAP_ adduct from 6-dEB. TE_DAP_ opened the 14-membered macrolactone ring, trapping the pyran form of the natural DEBS heptaketide as a stable acyl enzyme adduct. Deconvoluted mass spectra show the incorporation of the heptaketide at DAP3043.

In addition, we predicted that, like the wild type, TE_DAP_ could open cyclic macrolactone products (**Fig. S1B**), yielding linear TE_DAP_-polyketide adducts (**Figs. 2, S5C-E, S6C-E**). Thus, we tested four macrolactones (6-dEB^34^, narbonolide^32^, tylactone^35^ and 10-dml) using the same protocol as for the linear substrates. PikAIV TE_DAP_ was fully modified by narbonolide and partially modified by 10-dml, but we detected no modification using either 6-dEB or tylactone. In contrast, DEBSIII TE_DAP_ was fully modified by opening of the non-native narbonolide, modified at lower levels with the native 6-dEB macrolactone or tylactone, but was not modified by 10-dml. The lack of reactivity of PikAIV TE_DAP_ with 6-dEB and tylactone and of DEBSIII TE_DAP_ with 10-dml was not due to the weaker nucleophilicity of DAP relative to the native Ser. The TE_WT_ enzymes also had no detectable reactivity with these substrates. Upon ring opening, narbonolide, 10-dml, 6-dEB and tylactone can spontaneously convert to the corresponding pyran through hemiketalization and dehydration^9^. We detected both linear and pyran forms for the ring-opened adducts of narbonolide and 10-dml, but only the pyran forms of 6-dEB and the tylosin octaketide (**Figs. 2, S5C, S6C-D**).

Thus, PikAIV TE_DAP_ was highly selective for native polyketides, but DEBSIII TE_DAP_ exhibited a broader substrate tolerance (**Fig. 2A**). The reactivity of DEBSIII TE_DAP_ is consistent with early studies demonstrating that wild-type DEBSIII TE can form macrolactones smaller and larger than the natural 14-membered 6-dEB^21,36–38^.

### Structure of natural heptaketide adduct of PikAIV TE_DAP_

PikAIV and DEBSIII TE_DAP_ were screened in crystallization experiments, and crystals were obtained for PikAIV TE_DAP_. Following proteolytic cleavage of the N-terminal His_6_ tag, PikAIV TE_DAP_ crystallized in a form unrelated to that of the previously reported structures^z^. Crystallization conditions for the new form were optimized with PikAIV TE_WT_ and then applied to the TE_DAP_-heptaketide. Crystallization of PikAIV TE_DAP_-heptaketide followed incubation of TE_DAP_ with narbonolide, the natural pikromycin 14- membered macrolactone product, under conditions leading to complete modification with the heptaketide (+352.49 Da), as determined by intact protein LC/MS analysis (**Fig. 2A**). The PikAIV TE_DAP_-heptaketide yielded a 2.8-Å structure (**Table S1**). We also solved a 3.1-Å structure of PIKAIV TE_DAP-EOS-NBO_ from partially deprotected protein (**Table S1**).

The linear heptaketide adduct at DAP1196 is fully resolved in the active sites of both monomers in the TE_DAP_-heptaketide crystals with no indication of the pyran form, which may have been disfavored by the high pH (8.5) of crystallization (**Fig. 3A**). The heptaketide adopts a U-shaped conformation in the acyl cavity, with the distal hydroxyl (O13) ideally situated for attack at the acyl enzyme linkage, here a stable amide. Hydroxyl O13 is within ∼4 Å of the DAP amide analog of the TE_WT_-heptaketide ester. The natural heptaketide is positioned similarly to the phosphonate pentaketide mimic we previously trapped in the PikAIV TE active site (**Fig. 3B**)^15^.

**Figure 3.**
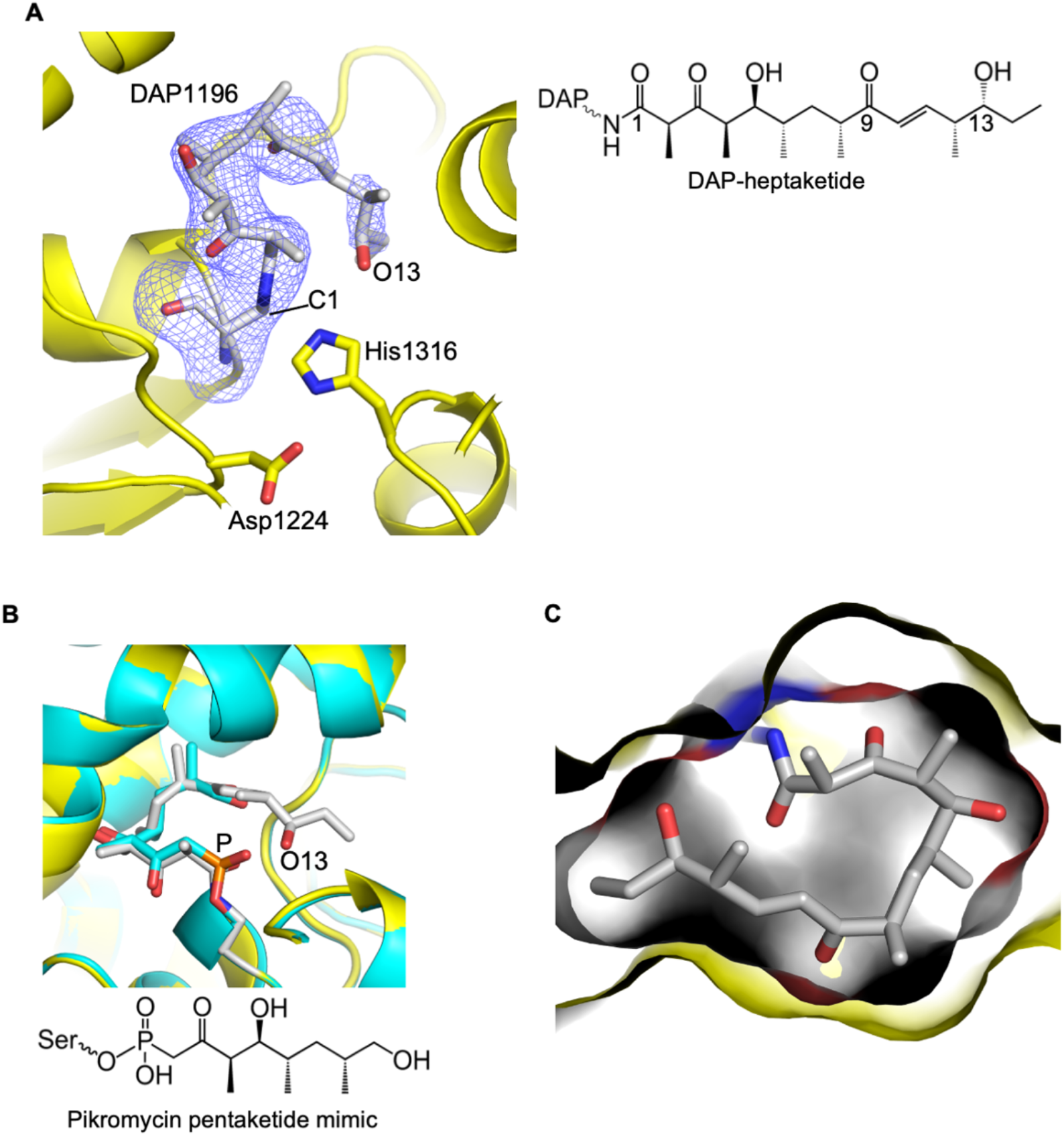
Crystal structure of PikAIV_DAP_-heptaketide. The heptaketide is curled in the acyl cavity with the substrate nucleophile (O13) near the acyl-enzyme linkage, here a stable amide. **A**. Electron density (3σ F_o_-F_c_ omit) for the heptaketide adduct at DAP1196. TE_DAP_ is rendered in cartoon form with the catalytic triad (DAP1196, Asp1224, His1316) as sticks with atomic coloring (white C for DAP-heptaketide, yellow C for TE, red O, blue N). **B**. Overlay of PikAIV TE_DAP_-heptaketide (yellow protein) with TE-pentaketide phosphonate mimic of the tetrahedral intermediate of acyl-enzyme formation (cyan protein, PDB 2HFJ). **C**. Shape complementarity of TE acyl cavity and heptaketide substrate. The image is a slice through the widest region of the acyl cavity (entrance from the left, exit to the right) illustrating the surfaces of enzyme and substrate.

The side chain hydroxyl of Thr1125 (helix α6) forms the sole hydrogen bond to the O9 carbonyl of both the heptaketide and the pentaketide mimic. The acyl cavity is filled with the heptaketide, with no space for water of hydrolysis. This contrasts with the pentaketide phosphonate analog, where the free end of the adduct exhibited greater flexibility (weaker electron density) within the acyl cavity^15^.

The acyl cavity was virtually unchanged by formation of the heptaketide adduct. Only a few side chains (Tyr1073, Leu1245, His1316, Phe1317) shifted slightly (<2 Å) from positions in the free enzyme to make space for the adduct. A remarkable hand-in-glove shape complementarity of the natural heptaketide and the acyl cavity (**Fig. 3C**) explains the high fidelity of macrolactone formation by the TE. This also explains the selectivity of PikAIV TE for natural substrates (**Fig. 2A, Fig. S5**). For example, the bulky Tyr1073 side chain would interfere with binding of the DEBS pathway heptaketide with its methyl substituent at C10. The tight fit also explains the stereoselectivity of the TE for the heptaketide O13 hydroxyl nucleophile as well as the inability of the tylosin octaketide, conformationally restricted by three double bonds in resonance, to occupy the active site. Contacts of the TE with the natural pikromycin heptaketide are predominantly hydrophobic (**Fig. S7**; Tyr1073 and Leu1077 in helix α2; Ala1126 in the TE core domain; Gln1231, Ile1234, Leu1241 and Leu1245 in helix α6; Met1261, Ala1265 and Leu1268 in helix α7; and Phe1317 adjacent to catalytic triad His1316). In contrast to these hydrophobic amino acids, the exit end of the TE tunnel is replete with polar side chains (Asp1071, Asp1078, Gln1231, Glu1235, Ser1238, Arg1266). These side chains form an effective “hydrophilic barrier” to an extended conformation of the of the heptaketide^15^.

### Acyl cavities in TE domains

For the PikAIV and DEBSIII TEs, several high-resolution crystal structures have been reported^13–15^, but these are lacking for the other three TEs in our panel. In order to compare acyl cavities, we solved crystal structures for the other three wild-type TEs, JuvEV TEWT, TylG5 TEWT and FluC TEWT (**Table S1**). These TEs possess the canonical α/β- hydrolase fold and other features common to PKS TE domains^13,15,17,18^ (**Fig. 4**). As in other offloading PKS TEs, two N-terminal α-helices form the dimer interface, and the catalytic triad is located at the center of a tunnel allowing ACP entry through one end and macrolactone or linear product formation in an acyl cavity at the opposite end. The JuvEV and TylG5 TE active sites contain a Ser-His-Asp catalytic triad (JuvEV Ser1637, Asp1664, His1756; TylG5 Ser1681, Asp1708, His1800), while the FluC TE has a rare Cys nucleophile (Cys2243, Asp2270, His2367).

**Figure 4.**
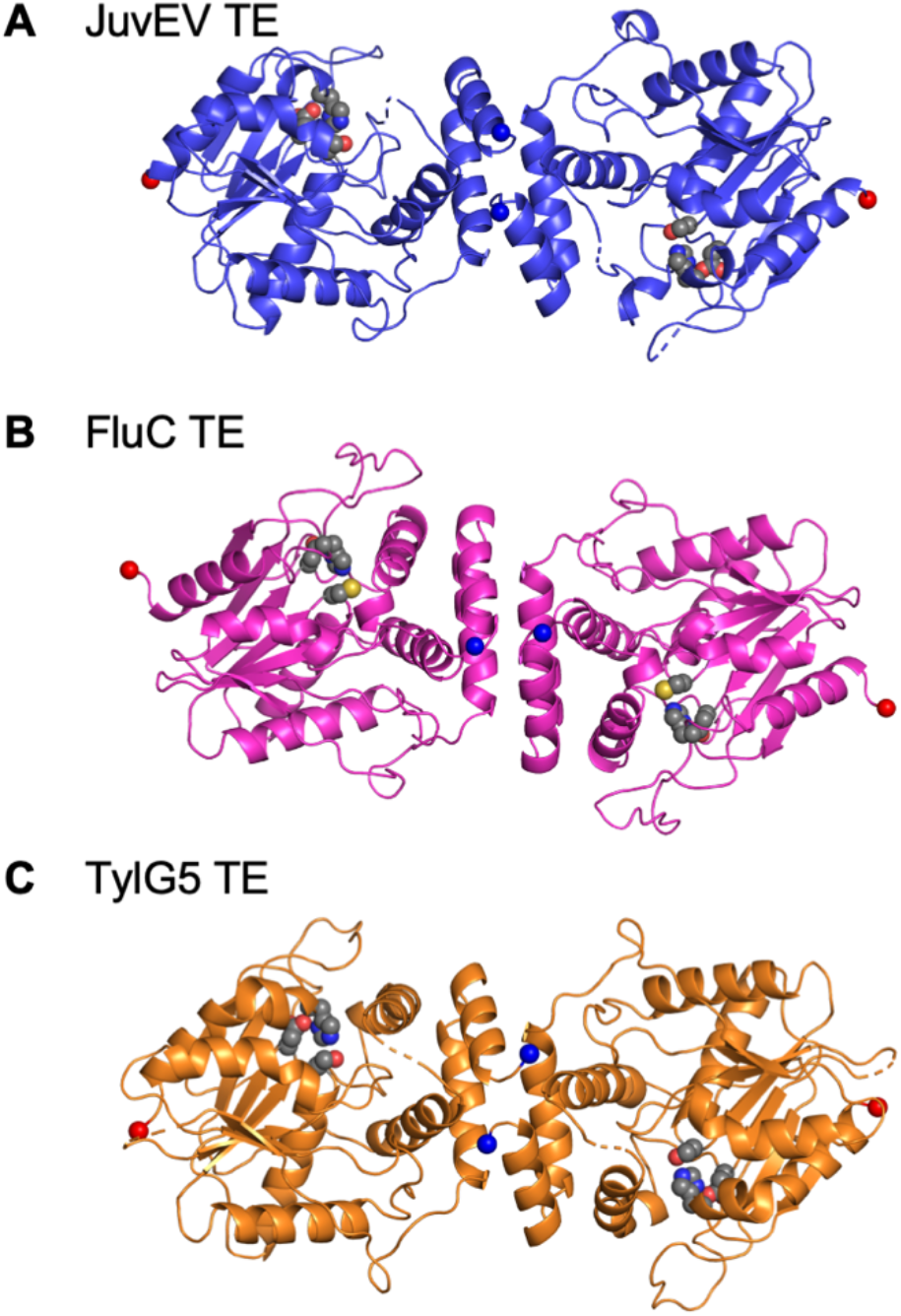
Structures of mPKS offloading TE domains determined in this study. **A**. JuvEV TE. **B**. FluC TE. **C**. TylG5 TE. Each TE is viewed along the dimer twofold with spheres indicating the polypeptide termini (blue N-terminus, red C-terminus). Amino acid side chains of each catalytic triad are shown as spheres (gray C, red O, blue N, yellow S).

We compared the acyl cavities of the seven offloading PKS TEs of known structure, including the five mentioned above and those from biosynthetic pathways for pimaricin (PimS4 TE)^17^ and tautomycetin (TmcB TE)^18^ (**Fig. 5**). Only the PikAIV and PimS4 TE crystal structures included a substrate or product. For the other cyclizing TEs, we positioned small-molecule crystal structures of the macrocyclic products^39–41^ in the acyl cavities of the corresponding TEs. Each acyl cavity has a shape and surface generally complementary to the product shape and surface, although the fits are not as snug as for the heptaketide trapped in PikAIV TE_DAP_. The interior surfaces of the acyl cavities are predominantly hydrophobic near the catalytic nucleophiles (Ser or Cys) and more polar at the tunnel exits. With the exception of the PimS4 TE, the macrocycle- forming TEs (**Fig. 5A-E**) have acyl cavities wide enough for the a linear polyketide to curl and bring the internal nucleophile near the acyl-enzyme linkage. Variability among the amino acid sequences and small differences in helix positions relative to the catalytic triad create slightly different shapes and orientations of the acyl cavities within these otherwise highly similar TE structures. The JuvEV and TylG5 TEs both produce tylactone (**Fig. 5C,D**), but a few sequence differences create somewhat different cavity shapes and, unlike the TylG5 TE, the exit half of the JuvEV TE acyl cavity has varying degrees of flexibility among the four independent views of the polypeptide in the crystal structure.

**Figure 5.**
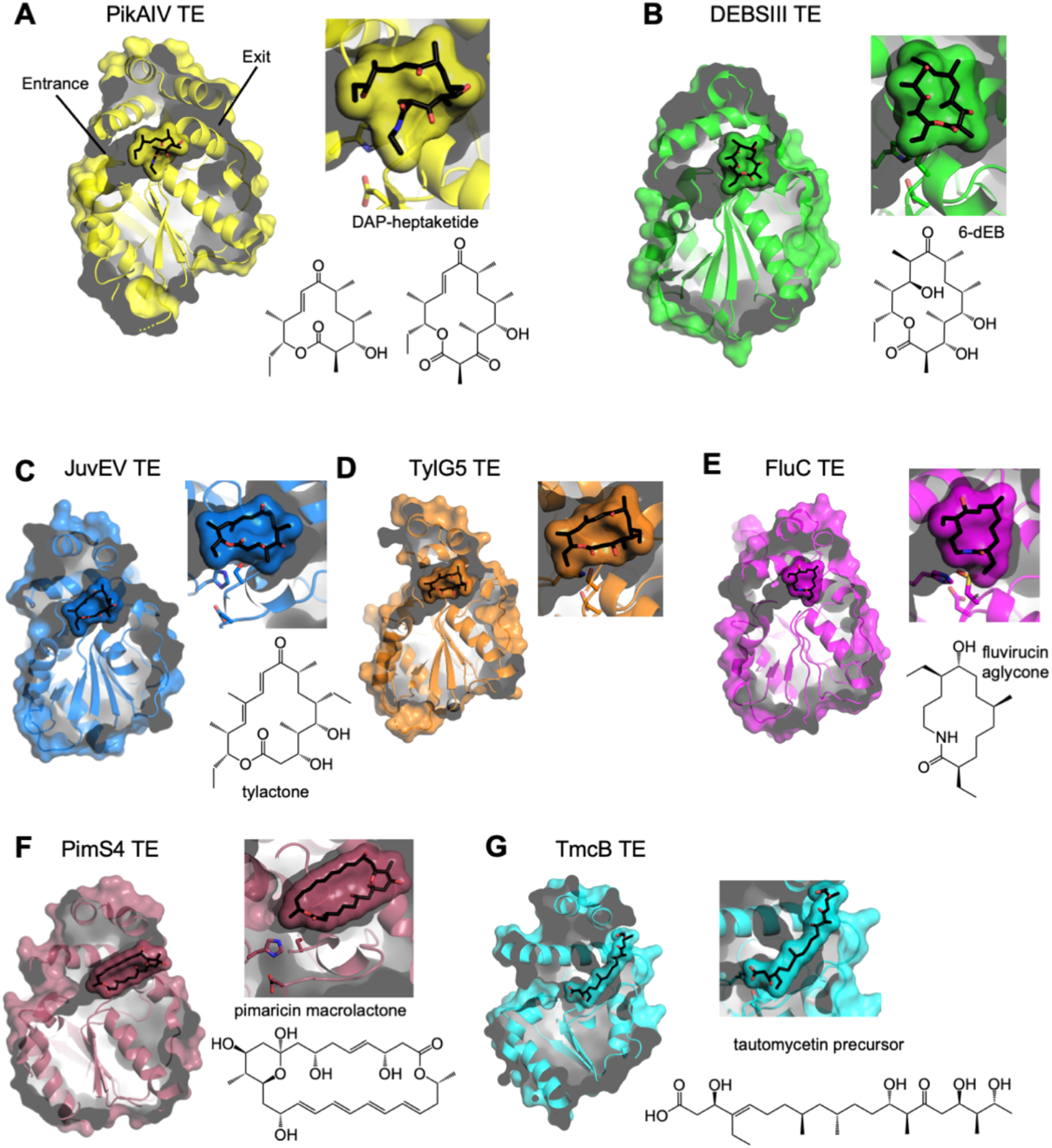
Acyl cavities in structures of mPKS offloading TEs. **A**. PikAIV TE_DAP_ (this study). **B**. DEBSIII TE (PDB 5D3K, modeled 6-dEB). **C**. JuvEV TE (this study, modeled tylactone). **D**. TylG5 TE (this study, modeled tylactone). **E**. FluC TE (this study, modeled fluvirucin aglycone). **F**. PimS4 TE (PDB 7VO5), a polyene TE. **G**. TmcB TE (PDB 3LCR, modeled product), which produces a linear polyketide. Cut-away views of the TE monomer structures show the active site tunnels in the viewing plane, with entrance from the left and exit to the right. The dimer interface is at the top. Acyl cavities are highlighted as semi- transparent surfaces Pik heptaketide in **A** and macrocyclic products in **B**-**F**. Products are adjacent to each TE; tylactone is the product of both JuvEV TE and TylG5 TE.

In contrast to the other cyclizing TEs, the PimS4 TE acyl cavity accommodates the 26- membered macrolactone product, but is too narrow for the polyene substrate to enter in an extended conformation and curl within the cavity (**Fig. 5F**). A pyranose ring at C13-C17 creates a bend in the substrate, suggesting it may enter the active site in a pre-curled form. The TmcB TE, the only TE in our panel that forms a linear product, has the narrowest acyl cavity and could not fit a curled substrate (**Fig. 5G**). Compared to the other PKS offloading TEs, the acyl cavities in the pimaricin and tautomycetin TEs are longer due to the absence of a bend at the start of helix α6 and narrower due to a shifted positions for helices α2 and α6. In the PimS4 TE, the product protrudes from the cavity exit, as would the undecaketide TmcB TE substrate.

## Discussion

Although first demonstrated with an NRPS TE, our study demonstrates the broad utility of the DAP unnatural amino acid to probe the substrate binding and selectivity of enzymes where a covalent intermediate forms at a side-chain hydroxyl (Ser, Thr) or thiol (Cys) nucleophile. The DAP incorporation reported herein enabled the first view of a PKS TE native-substrate acyl-enzyme. The structure of the near-natural catalytic intermediate of PikAIV TE provides mechanistic information about the critical cyclization step in macrolide antibiotic biosynthesis. The structure of the PikAIV TE_DAP_-heptaketide also illustrates potential challenges in developing/adapting biocatalysts to form macrolactones.

The natural linear heptaketide substrate in PikAIV TE_DAP_ is poised for macrolactone formation (**Fig. 3**). The substrate binds in a curled conformation that is highly complementary to the shape of the acyl cavity. This exquisite fit places the substrate nucleophile (O13 hydroxyl) in an ideal position for attack on the acyl enzyme linkage and subsequent macrolactone formation.

We found a variety of shapes and orientations for the acyl cavities in other offloading PKS TEs (**Fig. 5**), consistent with the sequence variability among amino acids lining the cavity and slight differences in helix positions relative to the catalytic triads. All the acyl cavities have a strongly hydrophobic surface in the region adjacent to the catalytic triad despite the variation in shape and orientation. In TEs that form 12-16-membered macrocycles (PikAIV, DEBSIII, TylG5, JuvEV, FluC), a bent helix α6 and a cluster of polar side chains at the acyl cavity exit induce the linear polyketide substrate to curl, bringing the internal nucleophile near the acyl-enzyme linkage (**Fig. 5A-E**). The cyclized products of these TEs fit within the non-polar region of the acyl cavities. Thus, we conclude that a “hydrophilic barrier” at the cavity exit^15^ assists catalysis where a flexible substrate must curl to position the internal nucleophile near the catalytic center. The shapes of the macrocyclic products also differ according to their double-bond content, substituents and chiral centers. Among the TEs that form 12-16-membered macrocycles, the seemingly subtle differences in acyl cavity shape must be well matched to the substrate to ensure efficient cyclization and avoid hydrolysis to the linear product. In contrast, the 26- membered polyene macrolactone product fills the acyl cavity and protrudes from the tunnel exit of the PimS4 TE. Here the pyranose ring may pre-form the substrate for cyclization (**Fig. 5F**)^17^. The tautomycetin TE produces a linear product and has an acyl cavity that is too slender for the polyketide to curl towards the acyl-enzyme linkage (**Fig. 5G**)^18^. The four hydroxyl substituents at the thioester-distal end of the TmcB TE polyketide are compatible with substrate extension into the polar exit end of the acyl cavity.

PikAIV TE can form a 14-membered macrolactone from the natural heptaketide intermediate as well as a 12-membered macrolactone from the natural hexaketide (**Fig. S1B**), both *in vitro* and in the *Streptomyces* producer^42^. The tight fit of natural substrate to acyl cavity may account for this rare dual-macrolactonization capability, but it also limits the substrate scope for macrolactone formation. The cavity shape appears unable to accommodate additional substituents (**Fig. 3C**), for example the methyl at C10 in the DEBSIII TE substrate (**Fig. S5D**).

Previous studies with variants of the natural hexaketide revealed an exquisite sensitivity to aspects of the substrate structure, leading to characterization of PikAIV TE as a gatekeeper for pathway fidelity^8^. Removal of methyl or ethyl substituents reduced the proportion of macrolactone relative to linear product, and inversion of either or both of the C12 (methyl substituent) and C13 (hydroxyl) chiral centers resulted in only linear products^8^. Reduction of the C9 carbonyl relaxes a conformational constraint (resonance of the carbonyl with the double bond at C10), but abolished macrolactone formation^20^.

In contrast to the PikAIV TE_DAP_, the DEBSIII TE_DAP_ had a broader substrate scope (**Fig. 2A**). DEBSIII TE_DAP_ reacted with the natural 6-dEB product and with non-natural macrolactones from the Pik and Tyl systems as well as Pik pathway intermediates, consistent with its reported production of macrocycles of various size^21,36–38^. Nevertheless, the DEBSIII TE reactivity is limited, as the linear product predominated if the chirality of the substrate nucleophile was inverted in a simplified unnatural substrate^23^. Additionally, we detected no ability of DEBSIII TE_WT_ or TE_DAP_ to catalyze rebound ring opening of the 12-membered macrolactone of the PikAIV TE product 10-dml (**Fig. S6E**).

TE formation of a macrolactone as opposed to a linear seco-acid product is the result of an internal substrate nucleophile out-competing water to resolve the acyl-enzyme. Poor positioning of the substrate nucleophile or substrate dynamics in the active site would disadvantage macrolactone formation. The observation of reduced or abolished macrolactone formation with non-natural substrates that are slightly misfit or more flexible in the acyl cavity highlights the kinetics of the competition with water – which nucleophile reaches the acyl-enzyme linkage first? In a previous study of the PikAIV TE^10^, substitution of the catalytic Ser with the stronger nucleophile Cys (TECys) increased the reaction rate (*k*cat) revealing a role for kinetics in reaction outcome^10^. Computed reaction trajectories for TE_Cys_ indicated an altered kinetic pathway compared to TE_WT_ . The increased *k*_cat_ of TE_Cys_ tipped the balance towards cyclization and away from hydrolysis for a non-natural stereodivergent (hexaketide C-11 OH) substrate where TE_WT_ catalyzed little or no cyclization^10,43^. The observation that greater speed promotes cyclization suggests that in the early enzyme-substrate complex the polyketide is curled in the active site with the substrate nucleophile poised exclusively for macrocyclization (outcompeting water).

Our study highlights the influence of acyl cavity shape on substrate selectivity and catalytic outcome for TEs that catalyze macrolactonization, thereby identifying the challenge of matching a substrate-enzyme pair to a desired macrolactone product. Structures of acylated forms of additional enzymes together with biochemical data and the use of a stronger Cys nucleophile could address/mitigate this challenge. This study is an important step towards realizing the promise of TE catalysts in development of improved macrolide antibiotics using native or engineered enzymes.

## Experimental Methods

### Construction of expression plasmids (Table S2)

DNA encoding JuvEV TE (amino acids 1508-1778, ARW71487) was subcloned from pET21b-JuvEV^35^ and inserted into pMCSG7^31^ using ligation independent cloning (LIC) to create expression plasmid pTMM19. PikAIV TE (1057-1346, AAC69332) was subcloned from pCAte2^20^ and inserted into pMCSG7 using Gibson assembly to create expression plasmid pTMM40. DNA encoding DEBSIII TE (2904-3167, AAV51822) was subcloned from pRSG33^44^ and inserted into pMCSG7 using Gibson assembly to create pTMM39. Synthetic DNA encoding the FluC TE (2106-2395, AFU66012)^45^ was inserted into pMCSG7 using Gibson assembly to create expression plasmid pTMM41. DNA encoding TylG5 TE (1549-1841, AAB66508) was subcloned from pDHS3007^46^ and inserted into pMCSG7 using Gibson assembly to create expression plasmid pTMM42.

For unnatural amino acid (UAA) incorporation, amber stop codons (UAG) were introduced into all TE expression plasmids in place of codons for the catalytic Ser or Cys residues using the QuikChange XL Site-Directed Mutagenesis kit (Agilent), optimized for GC-rich sequences (JuvEV TE S1637amb: pTMM47, PikAIV TE S1196amb: pTMM48, DEBSIII TE S3030amb: pTMM49, FluC TE C2243amb: pTMM50, TylG5 TE S1681amb: pTMM51). For small-scale optimization experiments with the UAA, sequences encoding all TE domains were amplified from amber codon-containing expression plasmids (pTMM47-51) and then inserted into pCDF7 (Novagen) vectors with identical construct boundaries (pTMM52-56), again using Gibson assembly. The pCDF7 expression vector encodes an N-terminal His_6_ tag but with a different selection marker (spectinomycin) and an origin of replication (p15A) compatible with the UAA system (pSF-DAPRS-PylT).

All primers are listed in **Table S3**. All constructs and mutations were verified by using the University of Michigan Sanger Sequencing Core, or by nanopore sequencing (Plasmidsaurus).

### Production of wild type proteins

All wild type TE domains were produced and purified in a similar manner. JuvEV TE, PikAIV TE and DEBSIII TE were purified in HEPES buffer. FluC TE and TylG5 TE were purified in Tris buffer. *Escherichia coli* BL21(DE3) cells were transformed with expression plasmids for PikAIV TE, DEBSIII TE or FluC TE. *E. coli* strain BL21(DE3) pRare2^47^ was transformed with the JuvEV TE plasmid and E. coli BL21(DE3) pGro7^48^ with the TylG5 TE plasmid. Cells were grown in 0.5 L of TB media with 100 μg/mL ampicillin to an OD_600_=1.5. Cultures producing JuvEV TE also contained 50 μg/mL spectinomycin (pRare2), and those producing TylG5 TE also contained 35 μg/mL chloramphenicol (pGro7). After reaching OD_600_=1.5, cultures were cooled to 20°C over 1 hr, induced with 200 μM IPTG and grown for 16 hr. Cultures producing JuvEV TE or TylG5 TE also had 1 g/L L-(+)-arabinose at induction. Cells were harvested by 30-min centrifugation at 8,000 rpm and cell pellets were stored at -20°C.

Cell pellets were resuspended in 35 mL lysis buffer (50 mM HEPES pH 7.4 or 50 mM Tris pH 7.4, 300 mM NaCl, 10% (v/v) glycerol, 20 mM imidazole, 0.1 mg mL^-1^ lysozyme, 0.05 mg mL^-1^ DNase, and 2 mM MgCl2), incubated on ice for 45 minutes, lysed by sonication, and centrifuged at 38,760 x g for 30 min at 4°C. The soluble fraction was filtered through a 0.45 μm syringe filter, loaded onto a 5 mL HisTrap column (Cytiva Life Sciences), washed with 10 column volumes of buffer A (50 mM HEPES pH 7.4 or 50 mM Tris pH 7.4, 300 mM NaCl, 10% (v/v) glycerol), and eluted with a linear gradient of 20-400 mM imidazole in buffer A.

For crystallization screening, the N-terminal His_6_ tags were removed from JuvEV TE, PikAIV TE and DEBSIII TE. For those samples, nickel affinity eluate containing enriched TE domain was pooled, augmented with tobacco etch virus (TEV) protease (1:30 protease:TE) in buffer B with 2 mM DTT, and dialyzed overnight at 4°C in buffer B (50 mM HEPES pH 7.4 or Tris pH 7.4, 150 mM NaCl, 10% (v/v) glycerol). Tag-free TEs were separated from any uncleaved TE and TEV protease by nickel affinity chromatography.

Following nickel affinity purification, all TEs were concentrated by filter centrifugation (Amicon) and further purified by size-exclusion chromatography (HiLoad 16/60 Superdex S75, Cytiva) in buffer B. All TE domains eluted at apparent molecular weights of 55-65 kDa, consistent with expected dimer molecular weights for each TE domain +/- the N-terminal His_6_-tag (∼3 kDa).

### DAP-EOS-NBO synthesis

The doubly protected form of the unnatural amino acid (2*S*)-2,3- diaminopropionic acid (DAP), (2S)-2-amino-3-{[(2-{[1-(6-nitrobenzo[d][1,3]dioxol-5- yl)ethyl]thio}ethoxy)carbonyl]amino}propanoic acid (DAP-EOS-NBO), was synthesized in house and by the Vahlteich Medicinal Chemistry Core (University of Michigan) using a published protocol_30_. The purity of the synthesized DAP-EOS-NBO was verified by LC/ESI-MS and ^1^H NMR (**Fig. S2**).

### DAP-EOS-NBO incorporation and TE_DAP_ purification

Extensive small-scale expression optimization was conducted to maximize incorporation of the DAP-EOS-NBO amino acid and TE_DAP_ production. *E. coli* BL21(DE3) cells were co-transformed with TE expression plasmids having an amber codon substituted for the catalytic Ser or Cys and with pSF-DAPRS-PylT^30^, which encodes an evolved *Methanosarcina barkeri* (Mb) pyrrolysyl-tRNA synthetase/suppressor tRNA pair for the DAP-EOS-NBO amino acid and was kindly provided by Jason Chin (MRC Laboratory of Molecular Biology). 5 mL cell cultures were grown in TB media in 50 mL conical tubes with 100 μg mL^-1^ ampicillin (pTMM47-51) or 50 μg mL^-1^ spectinomycin (pTMM52-56), 50 μg mL^-1^ kanamycin (pSF-DAPRS-PylT), and 1 mM DAP-EOS-NBO resuspended in 0.5 M NaOH. The cultures were grown in the dark at 37°C to OD600=1.2, cooled to 20°C over 1 hr, induced with 200 μM IPTG, and grown for 16 hr. Growth in the dark prevented premature photo- deprotection of the DAP-EOS-NBO. Cells were harvested by 30-min 4°C centrifugation at 4,000 rpm and cell pellets were stored at -20°C.

Harvested cells from 5 mL cultures were resuspended in 1 mL HEPES or Tris buffer A, lysed for 15 min at room temperature using CelLytic Express (Sigma) following the manufacturer’s instructions, and centrifuged for 15 min at 4°C and 14,000 rpm. The lysate supernatant was incubated with 50 μL Ni-NTA resin (ThermoFisher) pre-equilibrated in buffer A at 4°C for 2 hr.

Nickel resin was washed three times with 100 μL HEPES or Tris buffer A and bound protein was eluted with HEPES or Tris buffer A with 400 mM imidazole. Production of TE_DAP_ was evaluated using SDS-PAGE. Suitable amounts of DEBSIII TE_DAP_ were obtained from pTMM39 or pTMM54 expression, and PikAIV TE_DAP_ from pTMM40 or pTMM53 expression. Expression tests for other TE_DAP_ proteins (JuvEV, FluC, TylG5) did not yield sufficient protein for analysis.

For large-scale production of TE_DAP_ domains, *E. coli* BL21(DE3) cells were co-transformed with pSF-DAPRS-PylT and expression plasmids for PikAIV TE_DAP_ (pTMM53) or DEBSIII TE_DAP_ (pTMM39) and grown at 37°C in 0.25 L TB media with 50 μg mL_-1_ kanamycin, 100 μg mL^-1^ ampicillin (pTMM39) or 50 μg mL^-1^ spectinomycin (pTMM53). DAP-EOS-NBO (1 mM in 0.5 M NaOH) was added 2-3 hr into cell growth. Cells were grown to OD_600_=1.2, cooled to 20°C over 1 hr, induced with 200 μM IPTG, and grown for 16 hr. Flasks were wrapped in aluminum foil for culture growth and expression to prevent photo deprotection. Cells were harvested by 30-min centrifugation at 4°C and 8,000 rpm and cell pellets were stored at -20°C.

PikAIV TE_DAP_ and DEBSIII TE_DAP_ were purified as for the wild type proteins, with a few modifications. DEBSIII TE_DAP_ was purified in Tris buffer. For photo-deprotection, the eluate from Ni-affinity purification (TE_DAP-EOS-NBO_) was pooled in a 50 mL conical tube and irradiated with UV light (365 nm) for 4 min at room temperature to remove the photolabile protecting group. N-terminal His tags were removed as for the wild type proteins. In the final size-exclusion chromatography step, PikAIV TE_DAP_ (MW 31.1 kDa) eluted at apparent molecular weight ∼60 kDa, and DEBSIII TE_DAP_ (28.6 kDa) at ∼58 kDa (**Fig. S3)**. TE_DAP-EOS_ was incubated 1 week at 4°C in pH 8.0 buffer to promote spontaneous loss of the remaining protecting group, yielding TE_DAP_, and deprotection was confirmed by mass spectrometry (see below, **Fig. S4**).

### Formation of TE_DAP_ acyl-enzyme adducts

Pikromycin thiophenol-pentaketide was synthesized and used with PikAIII and PikAIV for in vitro biocatalysis of narbonolide as previously described^32^. Pikromycin C3-hydroxymethyl-thiophenol-hexaketide substrates was also synthesized as previously described^33^. Macrocyclic 6-deoxyerythronolide B (6-dEB)^34^ and tylactone^35^ were chemoenzymatically synthesized in vitro.

TE_DAP_-polyketide adducts were formed by 24-hr incubation at room temperature of a 5 μL reaction mixture (40 μM PikAIV TE_DAP_ or DEBSIII TE_DAP_, 160 μM polyketide substrates/products, 50 mM HEPES pH 8.0, 150 mM NaCl, 10% (v/v) glycerol). Reactions were initiated by addition of the polyketide substrate or product. Reaction mixtures were diluted to 4 μM TE_DAP_, centrifuged 15 min at 4°C and 13,000 rpm, and analyzed by LC-MS as described below.

### LC-MS analysis of TEs

An Agilent 6545 Q-TOF mass spectrometer operated in positive-ion mode with an Agilent 1290 HPLC system was used to assess the deprotection state of PikAIV TE_DAP_ and DEBSIII TE_DAP_. 2 μL of TE_DAP_ (4 μM) underwent reverse-phase HPLC (Agilent PLRP-S reversed phase column 3.0 μM, 50 x 2.10 mm) in H_2_O with 0.2% (v/v) formic acid at a flow rate of 0.2 mL min^-1^. Protein was eluted over a 8 min linear gradient of 5-100% acetonitrile with 0.2% (v/v) formic acid. MS conditions were as follows: fragmentor voltage, 225 V; skimmer voltage, 25 V; nozzle voltage, 1000 V; sheath gas temperature, 350°C; drying gas temperature, 325°C. Data were processed using the MassHunter BioConfirm 10.0 software (Agilent).

### Crystallography

Crystals of all TE domains were grown by sitting-drop vapor diffusion at 20°C. All crystallization conditions included a cryoprotectant, and crystals were flash cooled in liquid N_2_ directly from their growth solutions.

JuvEV TE (residues 1508-1778) crystals grew in a 1 μL:1 μL mixture of protein (∼10 mg/mL in HEPES buffer B without glycerol) and reservoir solution (20 mM MgCl_2_, 22% (w/v) polyacrylic acid 5100, 100 mM HEPES pH 7.5). Large, thin, rod-shaped crystals formed in 1 week.

FluC TE (residues 2106-2395) crystals grew in a 0.5 μL:0.5 μL mixture of protein (∼10 mg/mL in Tris buffer B) and reservoir solution (100 mM sodium acetate, 30% (v/v) PEG 400, 100 mM MES pH 6.5). Small hexagonal crystals grew within 2-4 days.

TylG5 TE (residues 1549-1841) crystals were grown using the sitting drop method at 20°C in a 1 μL:1 μL mixture of protein (∼10 mg/mL in Tris buffer B) and reservoir solution (1.0 M sodium citrate, 100 mM sodium malonate, 100 mM Bis-Tris propane pH 6.5). Crystals grew in clusters of hexagonal rods.

PikAIV TE_WT_ (residues 1057-1346), PikAIV TE_DAP_, TE_DAP-EOS-NBO_ and TE_DAP_-heptaketide crystallized under identical conditions. Crystals grew in a 0.5 μL:0.5 μL mixture of protein (15-20 mg/mL in HEPES buffer B) and reservoir solution (1.2 M LiCl, 25% (w/v) PEG 4000, 100 mM HEPES pH 8.5). Large rectangular (WT) or rod-shaped (DAP) crystals formed in 3-5 days.

All diffraction data were collected at 100 K on GM/CA beamline 23ID-B or 23ID-D at the Advanced Photon Source (APS) at Argonne National Laboratory (Argonne, IL) (**Table S1)**. Data were processed and scaled using XDS^49^, with the exception of the TylG5 TE data, which were indexed and integrated in MOSFLM^50^ and scaled in SCALA in the CCP4 suite^15^. All manual model building was done in Coot^51^.

The TylG5 TE yielded crystals apparently in Laue group 6/mmm, but the intensity statistics indicated twinning^52^. Automated molecular replacement in the BALBES pipeline^53^ detected twinning operator K, H, -L with a refined twin fraction (α) of 49%, and identified a solution in space group *P*6_5_ using 42% identical PikAIV TE^15^ (2HFJ) as a search model. Manual model building was iterated with refinement in REFMAC5^54^ and BUSTER^55^. TLS groups were manually defined based on groups proposed by the TLSMD server^56^.

The structure of JuvEV TE was solved by molecular replacement with Phaser^57^ in the Phenix^58^ software suite using the 63% identical TylG5 TE as a search model. Four copies of the TylG5 TE monomer were placed, and translational non-crystallographic symmetry (tNCS) was indicated by a peak at (0, ½, 0) of height 73.4% relative to the origin of the native Patterson map. The structure was completed by iterative rounds of model building in Coot and refinement in phenix.refine^59^. Electron density for helix *α*6 in the JuvEV TE active-site lid was poor. An AlphaFold2^60,61^ (AF2) prediction was an excellent match to the crystal structure for most of the

JuvEV TE, but helix *α*6 was predicted at low confidence (pLDDT 70-80). Nevertheless, the AF2 model had a reasonable fit to weak electron density for part of helix *α*6 in chains C and D. This helix was omitted from chains A and B where no density was visible.

The FluC TE (residues 2106-2395) crystal structure was solved by molecular replacement in Phaser using the 36% identical PikAIV TE (2H7X)^29^ as a molecular replacement search model and refined in phenix.refine.

With His tags removed, wild type PikAIV TE and all DAP variants crystallized isomorphously, but in a different form than for the published PikAIV TE structures. The structure of tag-free wild-type PikAIV TE was solved by molecular replacement with Phaser using our previous His-tagged PikAIV TE structure^15^ (2HFK). The new PikAIV TE_WT_ structure was then used to solve structures of PikAIV TE_DAP_, TE_DAP-EOS-NBO_ and TE_DAP_-heptaketide. For model refinement, stereochemical restraints for DAP-EOS-NBO and DAP-heptaketide were generated using eLBOW^62^, and the adducts were defined as L-amino acids in the protein. Subsequent model building was performed in Coot and refinement in phenix.refine.

Structures were validated in MolProbity^63^ (**Table S1**). All structure images were generated in PyMOL^64^, and sequences were aligned with Clustal^65^ in Jalview^66^.

## Supporting Information

For the panel of five TE domains, multiple sequence alignment and substrate/product structures; LC/MS and NMR validation of doubly protected DAP-EOS-NBO; purification of PikAIV TE_DAP_ and DEBSIII TE_DAP_; LC/MS validation of deprotection to yield PikAIV TE_DAP_ and DEBSIII TE_DAP_; PikAIV TE_DAP_ reaction with a substrate panel; DEBSIII TE_DAP_ reaction with a substrate panel; contacts of PikAIV acyl cavity with TE_DAP_-heptaketide; table of expression plasmids; table of PCR primers; crystallographic information table.

## Author Information Corresponding Author

Janet L. Smith – University of Michigan, Life Sciences Institute, Ann Arbor, Michigan 48109, United States; University of Michigan, Department of Biological Chemistry, Ann Arbor, Michigan 48109, United States; https://orcid.org/0000-0002-0664-9228;

## Authors

Tyler M. McCullough – University of Michigan, Life Sciences Institute, Ann Arbor, Michigan 48109, United States; University of Michigan, Department of Biological Chemistry, Ann

Arbor, Michigan 48109, United States; Present address: Clarion life science consultancy, Boston, Massachusetts, United States; https://orcid.org/0000-0002-5949-8777

Vishakha Choudhary – University of Michigan, Life Sciences Institute, Ann Arbor, Michigan 48109, United States; University of Michigan, Department of Biological Chemistry, Ann Arbor, Michigan 48109, United States; https://orcid.org/0000-0001-7722-1758

David L. Akey – University of Michigan, Life Sciences Institute, Ann Arbor, Michigan 48109, United States; Present address: University of Michigan, Department of Molecular, Cellular, Developmental Biology, Ann Arbor, Michigan 48109, United States; https://orcid.org/0000-0002-5687-540X

Meredith A. Skiba – University of Michigan, Life Sciences Institute, Ann Arbor, Michigan 48109, United States; University of Michigan, Department of Biological Chemistry, Ann Arbor, Michigan 48109, United States; Present address: Harvard Medical School, Department of Biological Chemistry and Molecular Pharmacology, Blavatnik Institute, Boston, Massachusetts 02115, United States; https://orcid.org/0000-0003-4615-6775

Steffen M. Bernard – University of Michigan, Life Sciences Institute, Ann Arbor, Michigan 48109, United States; University of Michigan, Program in Chemical Biology, Ann Arbor, Michigan 48109, United States; Present address: Vividion Therapeutics, San Diego, California 92121, United States; https://orcid.org/0000-0001-8061-6136

Jeffrey D. Kittendorf – University of Michigan, Life Sciences Institute, Ann Arbor, Michigan 48109, United States; Present address: Pharmaforensics Laboratories, Ann Arbor, Michigan 48103, United States; https://orcid.org/0009-0005-2799-680X

Jennifer J. Schmidt – University of Michigan, Life Sciences Institute, Ann Arbor, Michigan 48109, United States; Present address: Apertor Pharmaceuticals Inc., Alameda, California 94502, United States

David H. Sherman – University of Michigan, Life Sciences Institute, Ann Arbor, Michigan 48109, United States; University of Michigan, Department of Medicinal Chemistry, Ann Arbor, Michigan 48109, United States; University of Michigan, Department of Microbiology & Immunology, Ann Arbor, Michigan 48109, United States; University of Michigan, Department of Chemistry, Ann Arbor, Michigan 48109, United States; https://orcid.org/0000-0001-8334-3647

## Author Contributions

T.M.M. and J.L.S. conceived of the project; T.M.M. developed expression systems for TE_DAP_ proteins; T.M.M., V.C., S.M.B., M.A.S. and J.D.K. developed expression systems and purified and crystallized wild type TEs. T.M.M and V.C. purified and acylated TE_DAP_ proteins and performed LC-MS analysis; T.M.M., V.C., D.L.A., S.M.B and M.A.S. performed crystallographic analysis; J.J.S. synthesized doubly protected DAP; D.H.S. and J.L.S. supervised the work; T.M.M. and J.L.S. prepared the manuscript; all authors read and agreed with the manuscript.

## Acknowledgements

The work was supported by NIH grant R01 DK042303 and the Rita Willis Professorship to J.L.S. and by NIH grant R35 GM118101 and the Hans W. Vahlteich Professorship to D.H.S. V.C. was supported by NIH training grant T32 GM140223. We thank Jason Chin (MRC, Cambridge, UK) for pSF-DAPRS-PylT encoding the pyro-Lys tRNA synthetase, the University of Michigan Vahlteich Medicinal Chemistry Core for synthesis of doubly protected DAP, and Wendy Feng (University of Michigan Life Sciences Institute Mass Spectrometry Core) for assistance with MS analysis. GM/CA@APS has been funded by the National Cancer Institute (ACB-12002) and the National Institute of General Medical Sciences (AGM-12006, P30GM138396). This research used resources of the Advanced Photon Source, a U.S. Department of Energy (DOE) Office of Science User Facility operated for the DOE Office of Science by Argonne National Laboratory under Contract No. DE-AC02-06CH11357.

## Notes

The authors declare no competing financial interest.

## Notes

### Competing Interest Statement

The authors have declared no competing interest.

